# PyRAD: assembly of *de novo* RADseq loci for phylogenetic analyses

**DOI:** 10.1101/001081

**Authors:** Deren A. R. Eaton

## Abstract

Restriction-site associated genomic markers are a powerful tool for investigating evolutionary questions at the population level, but are limited in their utility at deeper phylogenetic scales where fewer orthologous loci are typically recovered across disparate taxa. While this limitation stems in part from mutations to restriction recognition sites that disrupt data generation, an alternative source of data loss comes from the failure to identify homology during bioinformatic analyses. Clustering methods that allow for lower similarity thresholds and the inclusion of indel variation will perform better at assembling RADseq loci at the phylogenetic scale.

*PyRAD* is a pipeline to assemble *de novo* RADseq loci with the aim of optimizing coverage across phylogenetic data sets. It utilizes a wrapper around an alignment-clustering algorithm which allows for indel variation within and between samples, as well as for incomplete overlap among reads (e.g., paired-end). Here I compare *PyRAD* with the program *Stacks* in their performance analyzing a simulated RADseq data set that includes indel variation. Indels disrupt clustering of homologous loci in *Stacks* but not in *PyRAD*, such that the latter recovers more shared loci across disparate taxa. I show through re-analysis of an empirical RADseq data set that indels are a common feature of such data, even at shallow phylogenetic scales. *PyRAD* utilizes parallel processing as well as an optional hierarchical clustering method which allow it to rapidly assemble phylogenetic data sets with hundreds of sampled individuals.

**Availability:** Software is written in Python and freely available at http://www.dereneaton.com/software/

**Supplement:** Scripts to completely reproduce all simulated and empirical analyses are available in the Supplementary Materials.

## 1 Introduction

Restriction site associated DNA libraries (RADseq; Baird *et al*., 2008) and related genotyping-by-sequencing approaches (e.g., GBS; Etter *et al*., 2011, ddRADseq; Peterson *et al*., 2012) take advantage of next-generation sequencing platforms to generate short reads from thousands of potentially homologous loci, across multiple individuals, by targeting genomic regions adjacent to restriction enzyme cut sites. In contrast to whole genome shotgun data, this provides a more efficient and economical source for performing comparisons across many sampled individuals, making it a popular tool for population genetic analyses (reviewed in Narum *et al*., 2013). The problem of efficiently obtaining sequence data across many individuals is also relevant to studies at deeper evolutionary time scales, such as genus or family level phylogenetics, and it is at this scale that RADseq is now receiving increased attention.

RAD sequences generated *in silico* from multi-species genome alignments have been shown to retain accurate phylogenetic information over evolutionary divergences as deep as 60 million years (Rubin *et al*., 2012; Cariou *et al*., 2013), however, to date all empirical RADseq studies conducted above the species-level were done at much shallower scales (The Heliconius Genome Consortium, 2012; Wagner *et al*., 2013; Wang *et al*., 2013; Eaton and Ree, 2013). Still, these studies confirmed theoretical expectations by demonstrating that concatenated, genome aligned, or even individual RADseq loci are capable of recovering well supported phylogenies and useful information about evolutionary discordance. Yet these studies also demonstrate that large amounts of missing data are a common feature of RADseq, which may limit its applications.

Theoretically, the ultimate scale over which RADseq data will be recovered, and thus phylogenetically useful, is inherently limited by mutations that disrupt restriction recognition sites, causing “locus dropout” – manifest as more missing data between more distantly related samples. However, empirically such limits are approached much sooner due to the loss of data to other technical factors related to data generation and analysis. For example, the choice of restriction enzyme, the size of selected fragments, and variation in sequencing coverage across samples all effect the number of loci recovered. Moreover, bioinformatic analyses used to assemble RADseq data sets recover different amounts of data depending on the parameters used to identify and cluster orthologs as well as filter for paralogs. In this paper I focus mainly on the latter aspect of locus dropout: factors limiting the identification (recovery) of orthologous loci when they are present in a data set.

Few software are currently available to assemble RADseq data (Chong *et al*., 2012; Peterson *et al*., 2012), the most commonly used of which is *Stacks* (Catchen *et al*., 2011, 2013). Here I describe a new software pipeline called *PyRAD*, which assembles orthologous RADseq loci into phylogenetic data sets with a focus on the analysis of highly divergent samples. The largest difference between *PyRAD* and *Stacks* is in their clustering algorithms used to identify homology among sequences. *Stacks* clusters reads by grouping those with less than N differences between them, not allowing for indels, whereas *PyRAD* uses a wrapper around the program *USEARCH* (Edgar, 2010), to instead measure sequence similarity by global alignment clustering, which incorporates indel variation. Below the *PyRAD* pipeline is described in detail, its performance is measured on simulated and empirical data, and it is compared with *Stacks*.

## 2 Methods

### 2.1 Summary of a *de novo* assembly

*PyRAD* requires as dependencies only the executable programs *USEARCH* (Edgar, 2010) and *MUSCLE* (Edgar, 2004), in addition to the common Python packages *Scipy* and *Numpy*. It is accessed through a command-line interface and employs a modular framework composed of seven sequential steps:

1. De-multiplexing (sorting by barcodes)
2. Quality filtering and removal of barcodes, cut sites, and adapters
3. Clustering within samples and alignment
4. Joint estimation of error rate and heterozygosity
5. Consensus base calling and paralog detection
6. Clustering across samples
7. Alignment across samples, filtering, and formatting

The de-multiplexing step (1) sorts raw FASTQ formatted sequence data into separate files for each sample, allowing a user set maximum number of mismatches (errors) in a barcode. If samples are already de-multiplexed this step can be skipped. Filtering (2) then removes barcodes, restriction recognition sites and Illumina adapters if present, and filters reads by their quality scores, replacing base calls below a user set limit with an ambiguous base (N). Reads with more than a user defined number of N’s are discarded. The clustering step (3) first collapses replicate sequences into individual records while retaining their total number of occurrences. Sequence order is randomized and clustering is performed using *USEARCH* with all heuristic options turned off. This creates clusters (stacks) by matching each sequential sequence to a ‘seed’ sequence that came before it, or else creating a new seed. The resulting stacks are aligned with *MUSCLE*.

The next step (4) uses the maximum likelihood equation of Lynch (2008) to jointly estimate the mean heterozygosity and sequencing error rate from the base frequencies at each site across all stacks in an individual (with greater than a set minimum depth of coverage), and uses these values (5) to calculate the binomial probability a site is homozygous (aa or bb) versus heterozygous (ab) (Li *et al*., 2008). A base call is only made if the depth of coverage is above a user set minimum, and high enough to make a statistical base call, else it is called undetermined (N). Consensus sequences containing more than a maximum number of undetermined sites are discarded. To filter for paralogs, consensus sequences are also discarded if they contain more than a maximum number of heterozygous sites, or more than the allowed number of haplotypes (two for diploids).

Consensus sequences are then clustered by sequence similarity across samples using USEARCH (6). The resulting stacks are aligned and filtered once again for paralogs before being output in a variety of formats (7). In this case, the number of samples over which any single site is heterozygous is used to identify potential paralogs under the assumption that polymorphisms are less likely to be retained over deep divergences than are fixed differences between paralogs. Step 7 can be repeated while substituting different subsets of taxa, and requiring different amounts of coverage across them, to construct data sets of varying size and completeness.

### 2.2 Hierarchical clustering for large data sets

As the size and scope of RADseq studies continues to grow the computational time required to assemble data sets can become burdensome. In *PyRAD* the limiting step for large data is often among-sample clustering (step 5), which is not parallelized. Clustering time scales with the total number of unique stacks and studies with many samples (*>*100) can quickly accumulate many low coverage or singleton loci that greatly slow run times. To speed this step an optional hierarchical clustering approach can be implemented which splits the job into smaller parallelized units. Here, loci are first clustered across samples within user-defined clades with each clade utilizing a separate processor. The seeds of these stacks are then used to cluster with seeds of stacks from other clades (Fig. 1), before a final stack is reconstructed from the matches to each seed from each hierarchical clustering step. A further gain in speed can be attained by excluding loci at each step that do not cluster with at least some minimum number of other samples, under the assumption that such loci are less likely to match with more distant relatives in the next step. Although this gain in speed comes at the cost of excluding loci with low taxon coverage, such loci are typically of little use in downstream analyses.

**Figure 1.**
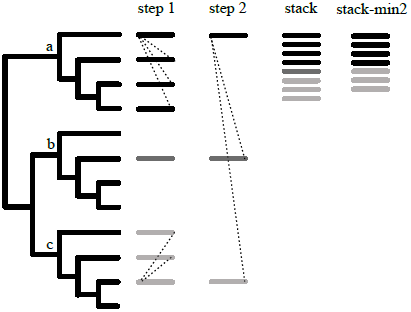
Optional hierarchical clustering approach for fast across-sample clustering of very large data sets in *PyRAD*. In the first iteration clustering is performed among samples within each user-defined clade (a, b, and c). Dotted lines show clustering of sequences to the randomly assigned seed sequence in each stack. In subsequent steps the seeds of stacks from previous steps are clustered with the seeds of stacks from other clades. Finally, matches to the seeds at each hierarchical step are reconstructed to build the full stack. A minimum cluster size can be set for each iteration to further increase speed at the cost of accuracy; in the example (min2) singleton loci are removed causing data from clade b to be lost.

### 2.3 Simulations

Sequence data were simulated on a species tree with 12 ingroup samples and one outgroup (Fig. 2) under a coalescent model within the Python package Egglib (Mita and Siol, 2012). Effective population size (50,000) and per base mutation rate (7×10^−9^) were constant across taxa, with the deepest divergence at 16 coalescent units. I simulated 10K loci for each of the 12 ingroup taxa at 20X coverage, with two alleles sampled from one diploid sample at 10X depth each. Reads were trimmed to 100 bp and a six bp barcode and five bp restriction recognition site were attached. Indels were introduced by sampling SNPs in the 12 ingroup taxa relative to the outgroup taxon at a probability of 0.0, 0.02, or 0.05, converting the derived allele into a deletion. This yielded three data sets that I refer to as the noIndel, lowIndel, and highIndel data sets.

**Figure 2.**
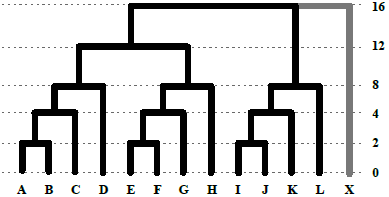
Species tree on which RADseq data were simulated under a coalescent model. Divergence times are in coalescent units. Mutations relative to the outgroup X were sampled at different frequencies to simulate deletions in three different data sets, and also to simulate disruption of the restriction recognition site in one data set.

A larger data set was also simulated on the same species tree to demonstrate the efficiency of the hierarchical clustering approach. In this case 500K loci were generated with no indels but allowing locus dropout through mutations that arise in a restriction recognition site relative to the outgroup. For this, I used a large region as the recognition site to create significant amounts of missing data between samples as expected in a deeply divergent phylogenetic data set. This led to a data set where each individual sample contained only approximately 150K loci, but for which the total data set comprised 500K loci potentially shared across samples. Thus the within-sample clustering step is relatively fast, whereas the across-sample clustering is much more computationally intensive, and benefits from the use of the hierarchical clustering. I refer to this as the noIndel-m data set.

All simulated data sets were compared using *PyRAD* v.1.621 and *Stacks* v.1.08. To the extent possible I used analagous parameters settings for both programs. For example, the clustering threshold of 85% in *PyRAD* is equivalent to 15 base differences in *Stacks* (parameters -m6 -M13-N15 -n15) given read lengths of 100 bp. A minimum stack depth of six was used in all analyses. Paralog filtering was not implemented on simulated data. The large noIndel-m data set was run with the same parameter settings, but was additionally run in *PyRAD* using a two-step hierarchical clustering approach. This first clustered loci within each of the three large four-taxon clades, then exluded any loci not shared across all four taxa within the clade, before subsequently clustering loci across clades.

### 2.4 Empirical data

A published RADseq data set from Eaton and Ree (2013) was downloaded from the NCBI sequence read archive (SRA072507). It includes data from 13 closely related taxa in the flowering plant genus *Pedicularis* sequenced with Illumina GAIIx, yielding 69 bp single end reads with barcodes and restriction sites removed. In contrast to the small simulated data sets which exhibit locus dropout due to bioinformatic errors alone, the empirical data have missing sequences owing to additional factors such as disruption to restriction sites and variation in sequencing coverage. It also contains low quality base calls and paralogs that must be filtered, making a comparison of *PyRAD* versus *Stacks* more difficult for this analysis.

To control for differences in how the two programs apply such filters, initial quality filtering was implemented only in *PyRAD*, with the resulting sequences analyzed by both programs. *Stacks* has fewer explicit filters to detect paralogs, but the closest approximations were used, and I report the results with and without paralog filters applied. The default parameters in *PyRAD* (maxHaplos=2, maxH=3, maxSharedH=3) were closely matched by using the following parameters, or manual edits, in *Stacks*, respectively: allowing only two haplotypes in a locus (–max_locus_stacks=2), removing loci from the final catalog if they contain more than four heterozygous sites, and removing stacks from the catalog if more than three samples are heterozygous for the same site. A clustering threshold of 85% was implemented in *PyRAD*, equivalent to 10 base differences in *Stacks* (parameters -m2 -M8 -N10 -n10) given the read lengths.

Detailed parameter settings and scripts to reproduce all simulated and empirical analyses are available as an IPython Notebook in the Supplementary Materials. Statistics measuring the distribution of taxon coverage across each assembled data set were measured from the results of each analysis found in either the “.haplotypes.tsv” output of *Stacks* or in the “.stats” output of *PyRAD*. Run times were also compared, where each analysis was run separately on a Linux machine with 24 3.47 GHz Intel Xeon processors, using 12 parallel processes in *PyRAD* or 12 threads in *Stacks*.

**Figure 3.**
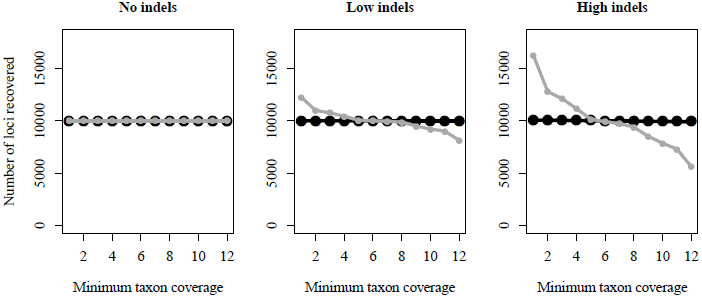
Comparison of *PyRAD* (black circles) versus *Stacks* (grey circles) in assembling simulated RADseq data sets that differ in their amount of indel variation. The two programs perform identically in the absence of indels (No indels), but in the presence of indels (Low indels & High indels) *Stacks* splits loci with indel variation into multiple loci with lower taxon coverage. Meaning a greater number of loci are shared across fewer taxa. *PyRAD*, in contrast, is little affected by indels, and recovers higher taxon coverage across all indel containing data sets.

## 3 Results

### 3.1 Simulation results

For the three small simulated data sets both *Stacks* and *PyRAD* retained all reads through the de-multiplexing and initial filtering steps. Similarly, within-sample clustering accurately recovered all 10K stacks for each taxon when either program analyzed the noIndels data set. In the lowIndels and highIndels data sets *PyRAD* performed nearly identically, recovering an average of 10000.0 and 10000.2 consensus sequences per sample, respectively, whereas *Stacks* recovered an average of 10027.4 and 10087.4 consensus sequences per sample; a result of splitting loci that are polymorphic for one or more indels within a sample.

The effect of indels is more pronounced when clustering across samples, where RAD sequences are more likely to exhibit indel variation. *PyRAD* recovered 10001, 10003, and 10086 loci in the three data sets with increasing proportions of indels, while *Stacks* recovered 10000, 12226, and 16285, respectively. Because only 10000 loci were simulated, those recovered beyond this number represent the splitting of loci caused by a failure to identify homology. Levels of sequence divergence are equal across the three data sets because indels were introduced by replacing mutations – meaning their phylogenetic distribution is identical to that of other mutations – thus the different results represent the effect of indel variation alone.

*PyRAD* identifies homology almost equally across the three data sets (Fig. 2), whereas *Stacks* performs worse when the data contain more indels, to the point of splitting nearly half the loci in the highIndels data set into multiple loci shared by fewer taxa. This led to lower average taxon coverage per locus in the assembled *Stacks* data sets, as well as fewer loci with full coverage across all taxa (Table 1). In all three small simulated data sets the *PyRAD* analysis finished in approximately half the time of *Stacks*.

#### 3.1.1 Large hierarchical clustering results

Both programs recovered similar results for the large noIndel-m data set in terms of average taxon coverage and number of loci shared across all taxa (Table 1). *Stacks* took over 25 hours to run while *PyRAD* finished in about half that time. Hierarchical clustering in *PyRAD* improved speed significantly, reducing the run time to less than three hours, but at the cost of reducing the average taxon coverage across the data set by one third. However, it had no effect on the percentage of loci recovered across all taxa, as the data that were excluded during hierarchical clustering represent loci shared by few samples. For very large data sets that include many sampled individuals and large numbers of loci the dramatic reduction in run times achieved through this approach will be useful.

**Table 1.** Comparison of *PyRAD* and *Stacks* in assembling simulated and empirical RADseq data sets of different sizes (Ntaxa & Nloci) and with different proportions of indels. Performance is measured as the average number of samples recovered per locus (Avg.cov) and percentage of Nloci with data recovered from all samples in Ntaxa (%Full.cov). The noIndel-m and empirical data sets experience locus dropout from mutations to restriction sites, and the latter also from variable quality and sequencing coverage. Run times (hours) are compared for assemblies utilizing 12 processors. Implementation of hierarchical clustering (*) with a minimum coverage of four reduced run times in *PyRAD*. Empirical results are shown with (-p) and without paralog filters applied, and the number of recovered loci (Nloci) was used to calculate %Full.cov.

| Data | Ntaxa | Nloci | Software | Avg.cov | %Full.cov | Time |
| --- | --- | --- | --- | --- | --- | --- |
| noIndel | 12 | 10K | *PyRAD* | 12.0 | 100.0 | 0.31 |
| noIndel | 12 | 10K | *Stacks* | 12.0 | 100.0 | 0.62 |
| lowIndel | 12 | 10K | *PyRAD* | 12.0 | 100.0 | 0.34 |
| lowIndel | 12 | 10K | *Stacks* | 9.8 | 81.5 | 0.65 |
| highIndel | 12 | 10K | *PyRAD* | 11.9 | 99.1 | 0.32 |
| highIndel | 12 | 10K | *Stacks* | 7.4 | 56.3 | 0.70 |
| noIndel-m | 12 | 500K | *PyRAD* | 4.5 | 0.1 | 10.47 |
| noIndel-m | 12 | 500K | *PyRAD* * | 2.9 | 0.1 | 2.95 |
| noIndel-m | 12 | 500K | *Stacks* | 4.5 | 0.1 | 25.70 |
| empirical | 13 | 178K | *PyRAD* | 3.0 | 1.4 | 36.42 |
| empirical | 13 | 207K | *Stacks* | 2.7 | 0.9 | 79.25 |
| empirical-p | 13 | 181K | *PyRAD* | 2.9 | 1.1 | 36.42 |
| empirical-p | 13 | 214K | *Stacks* | 2.4 | 0.5 | 79.25 |

### 3.2 Empirical results

When assembled by *PyRAD* the empirical data set recovered 181289 loci of which 56.4% were singletons, but for which more than 43K had taxon coverage of four or more, and 2465 were recovered across all taxa. In comparison *Stacks* recovered 213742 loci with a greater proportion of singletons (59.4%) and only 1940 loci shared across all taxa (Table 1). This difference is more striking after applying paralog filters to both analyses, wherein *PyRAD* recovers 1989 loci across all taxa but stacks recovers only 1123. Overall, *PyRAD* recovered more loci at high taxon coverage and fewer loci at low coverage than *Stacks* (Fig. 4).

**Figure 4.**
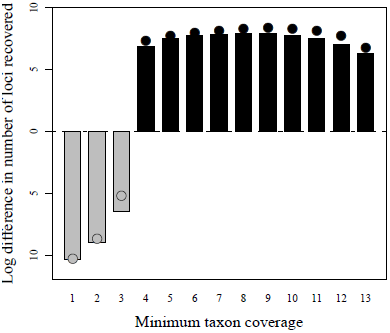
The log difference in number of loci recovered at each level of taxon coverage from the empirical RADseq data set of Eaton and Ree (2013) when analyzed by *PyRAD* and *Stacks*. Below the center line *Stacks* recovered more loci (grey), while above the line *PyRAD* recovered more (black). Circles indicate the comparison on data not filtered for paralogs and the bars after filtering for paralogs. *PyRAD* recovered more loci shared across a greater number of taxa, making its assembled data set more phylogenetically informative.

In the minimally phylogenetic informative data set (taxon covarage great than three) from the paralog filtered empirical *PyRAD* output, 32% of loci contained at least one indel, and of the loci with full taxon coverage, 15% contained indels. Comparing these values to the frequency of indels in the small simulated data sets, where 21% and 48% of loci contained indels (for the lowIndels and highIndels data sets, respectively) shows that the simulated values represent realistic proportions of indel variation as expected across a similar sized data set at a shallow phylogenetic scale.

## 4 Discussion

### 4.1 RADseq at the phylogenetic scale

One of the greatest barriers to RADseq phylogenetics stems from the significant amounts of missing data that are expected to result from mutations to restriction recognition sites. However, no empirical studies have yet shown whether this form of locus dropout is in fact the primary source of missing data. Instead, insufficient sequencing coverage and a failure to identify homology during bioinformatic analyses may account for a significant proportion of locus dropout, particularly in studies at shallow phylogenetic scales. Here I show that *PyRAD* performs better than the current *de facto* alternative software for assembling RADseq data sets from divergent samples that include realistic proportions of indel variation.

Indel variation is common even within a single species (Mullaney *et al*., 2010) and thus it should be no surprise that restriction site associated libraries often sample regions containing such variation. Within repetitive DNA elements (i.e., simple sequence repeats, microsatellites) indels can arise rapidly through slipped-strand mispairing during DNA replication (Levinson and Gutman, 1987), giving rise to single or often tandem repeat nucleotide differences between taxa. Rather than excluding or splitting loci that differ by short repeats, accurately clustering and aligning them across disparate species will help to create more complete and phylogenetically informative data sets.

### 4.2 Options and extensions

#### 4.2.1 Building contigs

The alignment clustering method used by *PyRAD* offers an added benefit through its ability to identify partial overlap among sequences. This proves particularly useful for GBS libraries that suffer from variable size selection which can result in overlap among paired-end reads, or between single-end reads sequenced from either end of short GBS fragments. Using reverse complement clustering *PyRAD* constructs contigs from overlapping regions with sufficient coverage to make consensus base calls. This allows building contigs longer than individual read lengths, and increases quality by combining sequences with high quality base reads at either end. Most importantly, it reduces duplication that would result from treating partially overlapping reads as separate loci.

#### 4.2.2 Introgression analyses

*PyRAD* is being developed to include a number of analysis tools in addition to data assembly. This currently includes measurement of D-statistics (Durand *et al*., 2011) and related tests for genomic introgression (Eaton and Ree, 2013). For this, *PyRAD* utilizes the distribution of RADseq markers as putatively unlinked loci, and performs non-parametric bootstrapping. A number of options are available to optimize the amount of shared data across samples when performing these tests, including SNP based tests and averaging allele counts across multiple sampled individuals.

## 5 Conclusion

*PyRAD* performs *de novo* assembly of RADseq data sets, including GBS, ddRADseq and paired-end reads. It is designed to perform equally well from population to phylogenetic scales, and is currently being implemented across this range, from linkage analyses among siblings to clade-level phylogenies with hundreds of tips. A major goal of this software was ease of use, and for this reason it utilizes common file formats familiar to those working in phylogenetics, and offers a variety of output formats for downstream analyses. Because the software does not need to be compiled *PyRAD* is easy to install on any desktop or computing cluster. It does not require large memory usage, and an analysis can be easily split into multiple shorter length jobs through its step-wise execution.

## Acknowledgement

Many thanks to R. Ree, B. Rubin and K. Ramanauskas for motivating the development of *PyRAD*, testing, and helping to identify bugs. Thanks also to R. Edgar whose software underlies much of the program, and for providing useful feedback.

### Funding

This research was supported by a National Science Foundation Dissertation Improvement Grant (DEB-1110598).

